# Non-visually-derived mental objects tax “visual” pointers

**DOI:** 10.1101/2025.05.02.651748

**Authors:** Xinchi Yu, Utku Türk, Allison Dods, Xing Tian, Ellen Lau

**Author notes:** **Corresponding author:** Xinchi Yu (;). Utku Türk and Allison Dods share second-authorship.

## Abstract

In daily life, we can construct mental objects from various modalities: we can create a cat in the mind from visual experience, or from what someone says or signs linguistically. Do mental objects derived from different input sources share the same set of “slots” (or pointers, indexicals)? Employing a novel imagine-one-then-more paradigm in a large EEG sample (*N* = 52) across two recording sites, we found that increasing the number of imagined objects derived from a non-visual source—auditory speech during eye closure—led to greater suppression of left posterior alpha power. This observation closely mirrors effects previously observed for visually-derived representations. Our results support the existence of a domain-general pointer system that functions as a modality-independent object register in moment-to-moment cognition.

## 1. Introduction

In our moment-to-moment cognition, we often need to keep one or more “objects” in the mind. These mental objects can be derived from a variety of sources, from direct visual input, to speech, to spontaneous thought (Brodbeck et al., 2016; Fukuda & Woodman, 2017). For example, the representation of a particular cat could be set up in the mind based on actually seeing a cat, or being told about a cat via spoken or sign languages. The most commonly studied scenario is when these objects are derived from direct visual input in recent history (i.e., within the same trial or experimental session). For example, in a trial of a typical visual working memory study (e.g. Luria et al., 2016), participants first see a memorandum consisting of one or more objects. After a short delay, they need to respond to a probe. To respond to whether the probe matches with the memorandum, one needs to hold the memorandum objects in mind across the delay. In this case, mental objects are derived from a direct visual source in recent history (i.e., the presented memorandum).

What about non-visually derived mental objects? Two opposing hypotheses reside in previous literature on cognitive science, including the philosophy of mind. Some propose that a difference in source suggests separate mental “slots” to represent these objects (Brody, 2020; Brody & Csibra, 2025; Murez et al., 2020), while others hypothesize that a common set of mental slots can be employed to represent objects derived from different sources (Brodbeck et al., 2016; Yu, 2024; Yu & Lau, 2023). According to the latter hypothesis, holding multiple cat individuals in the mind after viewing them or based on linguistic instructions (e.g., being told that “I saw a brown cat and a yellow cat yesterday in our yard”) would tax shared object-based resources in the mind, which would not be predicted by the former hypothesis. Of note, recent years have seen advances in understanding the nature of mental “slots”, particularly in the domain of visually-derived mental objects. Advancing from the somewhat vague conception of “slots”, recent theories propose that an object is mentally registered by a content-free pointer (or token, indexical), acting as a mental stand-in for this object and binding its features together (Awh & Vogel, 2025; Balaban et al., 2019; Ngiam, 2024; Swan & Wyble, 2014; Thyer et al., 2022; Wei et al., 2024; Yu & Lau, 2025). Critically, in this view, pointers are represented separately from features, corresponding to object-based load and feature-based load respectively. Empirically, object-based load can be isolated by controlling for the amount of features (Balaban & Luria, 2017; Naughtin et al., 2016). For example, four objects with the same color are expected to tax comparable feature-based load as one object with this color, while differing in object-based load, i.e., the number of pointers taxed (Gao et al., 2011; Naughtin et al., 2016; Xu, 2009).

Accompanying this theoretical trend, recent years have seen a surge in interest in exploring neural markers tracking the number of pointers in moment-to-moment processing (Jones, Diaz, et al., 2024; Jones, Thyer, et al., 2024; Thyer et al., 2022; Yu, 2024). For example, an evoked response—the negative slow wave (NSW)—appears to track visual working memory load (Feldmann-Wüstefeld, 2021; Fukuda et al., 2015; Yu, Li, et al., 2024). However, a trial in the current mental imagery study necessarily takes multiple seconds, and evoked sustained responses over longer EEG epochs are easily derailed by much larger yet non-neural drifts (Kappenman & Luck, 2010; Luck, 2014). In the current study, we instead examine posterior alpha power (8-12 Hz). Analysis of the alpha band is unlikely to be affected by slow drifts at multi-second level, and alpha power is also known to track visual working memory load (Diaz et al., 2021; Fukuda et al., 2015; Fukuda & Woodman, 2017). Encouragingly, posterior alpha power tracks object-based load when the number of features and the spatial extent of attention are controlled for (Chen et al., 2024). This allows us to use posterior alpha power as a measure of the number of pointers a participant is currently holding in mind.

Unlike existing studies with visual stimuli, in the current study, we examine a non-visual scenario—using verbal instructions to generate visual imagery—and we test whether object-based load in this case modulates posterior alpha power in a similar fashion to visually-derived object load. Specifically, we asked participants to imagine objects (identical in features) in the mind based on auditory speech instructions during eye closure. These objects were not visually presented to the participants at all (**Figure 1A**). Intriguingly, we found that speech-induced mental object load also suppresses posterior alpha power, similar to what was previously reported for visually-induced mental object load. This suggests that both visually and non-visually induced objects rely on a shared set of pointers in the mind.

**Figure 1.**
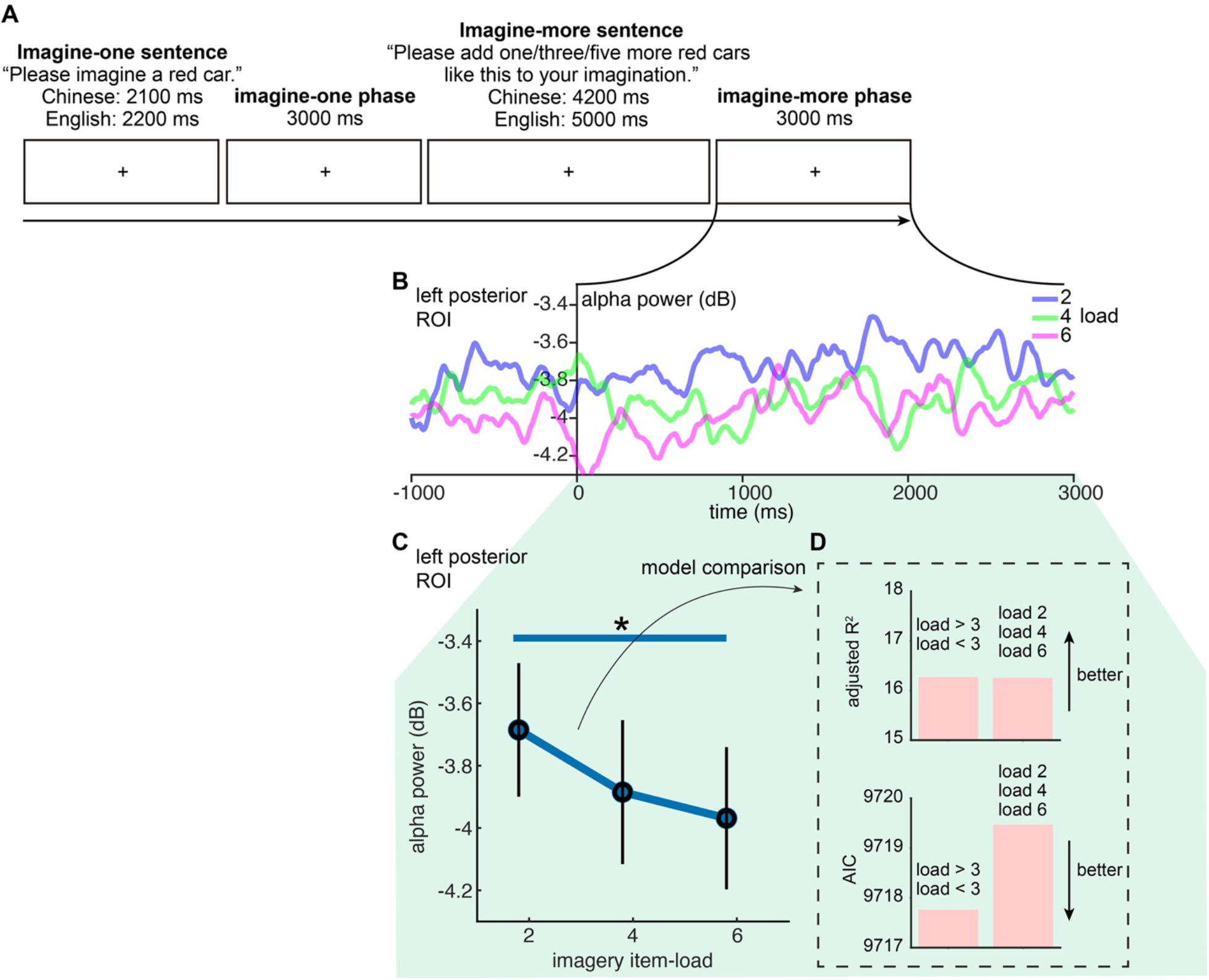
Left posterior alpha power inversely predicts speech-induced mental imagery load. (A) Illustration of the procedure of a trial in our experiment. (B) Average alpha power during the imagine-more phase and the preceding 1000 ms in the left posterior ROI. Data were all resampled to 500 Hz and smoothed with a sliding 100 ms window for visualization. For the data pattern across the entire epoch see **Supplementary Figure S1**. (C) Average alpha power across the imagine-more phase for the left posterior ROI. Error bars represent standard error across participants. *: *p* < 0.05. (D) Model comparison for the left ROI, between a GLME model with three load levels and one with only two load levels (load < and > 3), on adjusted R^2^ and AIC.

## 2. Materials and Methods

### 2.1. Participants

The entire dataset consists of 52 participants (age 18-29, 31 female), half tested in Shanghai, and half tested in Maryland. The sample sizes for each testing site were identical, and are comparable to previous studies on posterior alpha power for visually-induced object load (Fukuda et al., 2015; Fukuda & Woodman, 2017). The participants received monetary reimbursement or course credits for their participation. Two additional participants were excluded from analysis because of not completing all trials (*N* = 1) or because a reference electrode became dislodged during the recording (*N* = 1). Participants in Shanghai were administered stimuli in Mandarin Chinese, and participants in Maryland were administered stimuli in English; they were all proficient in the corresponding language. The procedures are approved by the IRBs of New York University Shanghai and University of Maryland College Park.

### 2.2. Stimuli

For each language (Mandarin Chinese and English), we constructed 28 adjective-noun phrases by crossing 6 monosyllabic color adjectives with 5 monosyllabic concrete nouns. 2 phrases out of the 30 combinations were excluded for each language for semantic ambiguities (e.g., “red shirt” may also mean “non-essential”). The adjectives and nouns for Chinese are hong2 (red); lü4 (green); huang2 (yellow); lan2 (blue); bai2 (white); hei1 (black); hua1 (flower); zhi3 (paper); bi3 (pen); che1 (car); wan3 (bowl). The adjectives and nouns for English are black, blue, gray, green, red, white, car, mug, shirt, spoon, bowl. The two lists are necessarily not exactly the same in meaning as some words are only monosyllabic in one language.

For each phrase, we generated an “imagine-one” sentence in the following form: *Please imagine a [adjective] [noun]* (Chinese: **请想象⼀** [classifier] [adjective] [noun]). The imagine-one sentence spans 2100 ms in Chinese and 2200 ms in English. For each phrase, we also generated an “imagine-more” sentence in the following form: *Please add [one/three/five] more [adjective] [noun] like this to your imagination* (Chinese: **请在想象中增加** [one/three/five] [classifier] 这样的 [adjective] [noun]). The imagine-more sentence had a duration of 4200 ms in Chinese and 5000 ms in English, and the number word had an onset at 2100 ms for Chinese and 800 ms for English. The sentences were synthesized with built-in Mac OS voice with consistent speech rate and syllable durations (Chinese voice: “Ting-Ting”; English voice: “Alex”). Each synthesized segment was normalized to the same intensity in Praat (Boersma, 2001).

### 2.3. Procedure

In each trial, the participants first saw a fixation cross for 1000 ms. Then they saw an instruction of “close your eyes then press space” on the screen, where they were instructed to first close their eyes, and subsequently press the spacebar to start a trial. After 1200 ms (Shanghai) or 8200 ms (Maryland), they heard one of the imagine-one sentences, e.g., “Please imagine a red car”. We chose to extend the time period between eye closing and onset of imagine instructions for the Maryland dataset because of recent observations that posterior alpha power may take several seconds to plateau after eye closure (Geller et al., 2014; Wöstmann et al., 2020). The participants were given a 3000 ms period to continue imagining the object, which we term the ‘imagine-one phase’. Then, they heard one of the imagine-more sentences with the same adjective-noun combination, e.g., “Please add three more red cars like this to your imagery”. This imagine-one-then-more procedure ensures that each imagined object was comparable in size across the three load conditions, mirroring the case of typical visual working memory experiments (Fukuda et al., 2015; Fukuda & Woodman, 2017; Xu & Chun, 2006). Following the imagine-more instruction, participants were given a 3000 ms period to continue imagining the object, which we term the ‘imagine-more phase’. Participants were explicitly instructed at the start of the experiment to imagine multiple objects *simultaneously* in the mind. Afterwards, they heard the command “stop”, and they were instructed to open their eyes and stop imagining. A self-paced inter-trial interval then followed before the start of the next trial. In line with the 84 imagine-more sentences, each participant went through 84 trials accordingly in random order. Prior to the main experiment, they were administered three practice trials with the adjective-noun combination of “green birds” (not present in the main experiment) to familiarize them with the procedure. The participants were also familiarized with the experimental word list beforehand to ensure they had no trouble comprehending the synthesized speech. Auditory stimuli were delivered binaurally via earphones for the Shanghai participants, and via a loudspeaker placed centrally in front of the Maryland participants.

### 2.4. EEG recording

#### Shanghai site

EEG was recorded from 32 electrodes mounted in an electronic cap (EasyCap; Brain Vision actiCHamp; Brain Products) according to the 10/20 system (FP1, Fz, F3, F7, FT9, FC5, FC1, C3, T7, TP9, CP5, CP1, Pz, P3, P7, O1, Oz, O2, P4, P8, TP10, CP6, CP2, C4, T8, FT10, FC6, FC2, F4, F8, FP2, Cz) at a sampling rate of 1000 Hz. Two additional EOG electrodes (HEOG and VEOG) were attached for monitoring ocular activity. The data were referenced online to electrode Cz. Impedances were maintained below 25 kΩ. The EEG signals were filtered online between DC and 200 Hz with a notch filter at 50 Hz.

#### Maryland site

EEG was recorded from 29 electrodes mounted in an electronic cap (Electro-Cap International Incorporated; SynAmps; Neuroscan) according to the 10/20 system (FP1, FP2, F7, F3, FZ, F4, F8, FT7, FC3, FCZ, FC4, FT8, T7, C3, CZ, C4, T8, TP7, CP3, CPZ, CP4, TP8, P7, P3, PZ, P4, P8, O1, O2) at a sampling rate of 500 Hz, with an additional electrode placed anterior to Fz as ground. Bipolar HEOG and VEOG electrodes were attached for monitoring ocular activity The data were referenced online to the right mastoid. Impedances were maintained at less than 25 kΩ for all scalp electrode sites, and less than 10 kΩ for mastoid and ocular electrodes. The EEG signals were filtered online between 0.05 and 100 Hz.

### 2.5. EEG data processing

EEG data were analyzed with customized MATLAB scripts (with EEGLAB and ERPLAB). Bad channels, if any, were interpolated spherically (0-1 for each participant, mean = 0.05). After removing DC bias, raw data from the scalp electrodes were re-referenced to their average, and then passed through a 0.1-40 Hz (cutoff frequencies) Butterworth filter. After that, ICA was conducted with the runica algorithm, and components that were labeled with “Muscle,” “Eye,” “Heart,” “Line Noise,” or “Channel Noise” with a confidence higher than 90% were removed from data, based on ICLabel (Pion-Tonachini et al., 2019). On average 0.4 components were removed for each participant.

To extract alpha power activities, data was subsequently bandpass-filtered with cutoff frequencies of 8 and 12 Hz with a Butterworth filter. Alpha power for each time point and electrode was calculated by squaring the analytic signal derived from Hilbert transformation (Adam et al., 2015; Foster et al., 2015). Epochs were extracted from 1000 ms before the offset of imagine-one sentences, until the end of the imagine-more phase. A very small proportion of epochs (0.27%) containing incidental presentation-timing fluctuations were excluded from the Chinese dataset. For each participant, outlier trials were excluded in case any electrode had a range more than 3 standard deviations from the mean range of that electrode over all trials. This resulted in 1,763 remaining (81% of all trials; two: 578; four: 597; six: 588) epochs in the Chinese dataset and 1,847 remaining (85% of all trials; two: 600; four: 625; six: 622) epochs in the English dataset. To facilitate comparisons across participants, alpha power in arbitrary units was then converted to decibels (dB) relative to the mean amplitude of the 1000 ms preceding the imagine-one phase (de Vries et al., 2017; Zeng et al., 2024). Guided by prior studies suggesting hemispheric differences in alpha power during mental imagery (Benedek et al., 2014; Moore & Lux, 1987), we treated the left (O1, P7) and right (O2, P8) posterior channels as separate ROIs. The time window of interest was the imagine-more phase, the imagination period immediately following the offset of the imagine-more sentence. Posterior alpha power was calculated by averaging the responses within the time windows and ROI channels for each trial.

### 2.6. Statistical analyses

To quantify the effect of load (2, 4 and 6) and ROI (left, right) on posterior alpha power during the imagine-more phase, we employed a generalized linear mixed-effect (GLME) model on trial-level data, with load, ROI and their interaction as fixed effects. Trial number was included as a random effect along with participant ID and recording site (Shanghai or Maryland) in this first model in order to account for trial-level relationships between the left and right ROI measurements: *Posterior alpha power ∼ 1 + load + ROI + load×ROI + (1*|*participant:site:trial)*. The response variable was modeled with a normal distribution and linked with an identity function.

After establishing the interaction effect between load and ROI, we then analyzed the data for the left and right ROIs separately. As there was only one value in each trial, in this model we included only participant and recording site as random effects: *Posterior alpha power ∼ 1 + load + (1*|*participant:site)*. Load and alpha power were z-scored prior to analysis. To complementarily examine “pairwise differences” between load conditions, we administered complementary GLME analyses, where load was instead coded as dummy variables: (a) two variables representing the effect of load 4 relative to 2, and load 6 relative to 2, or (b) two variables representing the effect of load 2 relative to 6, and load 4 relative to 6.

In order to evaluate whether there was a plateau beyond ∼ 3 items for the left ROI, we compared across two GLME models: one with three categorical load levels, and one with only a distinction between load < 3 and load > 3. The models were compared with a log-likelihood ratio test, and also on AIC (Akaike information criterion) and adjusted R^2^ (cf. Yu, Thakurdesai, et al., 2024). All reported *p*-values are two-tailed.

## 3. Results

GLME analysis revealed a significant interaction effect between ROI (left, right) and load: *β* = 0.023, SE = 0.011, *t* = 2.13, *p* = 0.033. Based on this interaction effect, we then analyzed the left and right ROIs separately. A significant effect of load was found in the left ROI, *β* = -0.031, SE = 0.015, *t* = -2.01, *p* = 0.044, but not in the right ROI, *β* = -0.007, SE = 0.015, *t* = - 0.46, *p* = 0.64 (**Figure 1B, C**). This lateralization effect was consistent with the visual impression from scalp distributions (**Supplementary Figure S2**). Complementary GLME analysis on the left ROI revealed a significant difference between load 2 and 6 (*β* = -0.075, SE = 0.037, *t* = - 2.01, *p* = 0.044), and a numerical trend between load 2 and 4 (*β* = -0.055, SE = 0.037, *t* = -1.48, *p* = 0.14). Coding the dummy variable instead relative to the load 6 condition, we similarly found a significant difference between load 2 and 6 (*β* = 0.043, SE = 0.022, *t* = 2.01, *p* = 0.044), but not between load 4 and 6 (*β* = -0.012, SE = 0.021, *t* = -0.54, *p* = 0.58).

A GLME model with three load levels (adjusted R^2^ = 16.25%, AIC = 9719.5) did not out-perform a similar model only with two load levels (with only a distinction above and below 3, adjusted R^2^ = 16.27%, AIC = 9717.8), mirroring the results of the log-likelihood ratio test, χ^2^_(1)_ = 0.29, *p* = 0.59. Rather, the three-load-level model had numerically lower adjusted R^2^ and higher AIC (**Figure 1D**). Taken together, modeling load conditions as three levels appeared not to be necessary, compared to the two-load-level model.

## 4. Discussion

### 4.1. Left posterior alpha power reflects object-based mental object load

With a novel imagine-one-then-more paradigm leveraging 3,610 EEG trials from 52 participants, we observed that left posterior alpha power decreases as object-based imagery load increases, when these mental objects were derived from a non-visual source: auditory speech instructions for imagery during eye closure. This load effect was object-based rather than feature-based, since participants were asked to imagine identical objects with shared features. Moreover, the alpha power appeared to plateau beyond ∼ 3 items. This load-dependent and capacity-limited pattern is in line with what has been observed for visually-induced object-based load (Chen et al., 2024; Fukuda et al., 2015; Fukuda & Woodman, 2017).

Intriguingly, our current EEG effect was left-lateralized. Indeed, the laterality of mental imagery has long been debated (Brown & Kosslyn, 1993; Ehrlichman & Barrett, 1983; Liu et al., 2022). Interest in this topic traces back to earlier neuropsychological investigations with e.g., split-brain patients whose corpus callosum was removed. For example, Farah and colleagues (1985) tested one such patient on a task where they would see an uppercase letter, and respond to whether the corresponding lower case letter only occupied the “middle” vertical range (e.g., a, c, e) or extended into upper or lower spaces (e.g., b, d, y). This task was believed to reflect mental imagery as one would need to imagine the exact shape of the lowercase letters. Strikingly, the patient could only perform such task above chance when the uppercase letter was presented in the right visual field, corresponding to the left hemisphere. In contrast, other studies suggest a primary role of the right hemisphere for face imagery (Bowers et al., 1991). As mental imagery is a multi-component construct consisting of at least the representation of pointers and features, as well as temporally distinct phases such as formation and maintenance (Farah, 1984; Kosslyn & Shwartz, 1977), the answer to the laterality question is unlikely simple. In other words, it is unlikely that mental imagery *only* relies on the left or right hemisphere. The exact brain regions recruited and their laterality may depend on a range of factors including the content that is imagined (e.g., objects vs. faces, Liu et al., 2022) and the cognitive computations required by the task (Hu & Yu, 2023; Kosslyn, 1988). While our current study was not designed to adjudicate this long-standing debate, we observed that imagery load for objects induced a left-lateralized EEG response. Given the limited spatial resolution of EEG, future studies employing neuroimaging methods with higher spatial resolution (e.g., MEG) will be critical in clarifying the laterality of neural sources for this effect.

### 4.2. The capacity of mental imagery

Our current results suggest that posterior alpha power plateaued beyond ∼ 3 items, in line with the well-established object-based capacity limits for visual working memory (∼ 3 items; Cowan, 2001; Awh et al., 2007; Zhang and Luck, 2008; Bengson and Luck, 2016). This is similarly consistent with the hypothesis that mental imagery makes use of the same object-based pointer system as standard visual working memory.

Interestingly, a recent study by Balaban and Ullman (2024) argued that participants could only simulate gravity-constrained movements of obscured visual items serially, even comparing two items to one item. In that study, participants needed to respond to when balls – based on gravity – would fall onto the ground, where only the initial part of the trajectories was shown. This means that the participants needed to mentally continue the trajectories temporally. For one ball, participants’ temporal estimate aligned linearly with the actual timing. However, when two balls were presented, although participants’ estimate for each item correlated well with the actual timing, one of the balls was consistently estimated as falling late – by about 600 ms. The authors took this to reflect a serial process instead of a parallel simulation process, implying a one-item capacity in this task. This intriguing result poses a seeming discrepancy with our current results, as a capacity limit of one item was suggested, instead of ∼ 3. We hypothesize that the one-item capacity limit observed in Balaban and Ullman (2024) arose from the communication bottleneck between object-based representations and the intuitive physics system. In our current task, participants only needed to represent items in the mind, while these items do not need to partake in computations such as the ones related to intuitive physics. Taken together, our findings and those of Balaban and Ullman (2024) call for future research to further characterize the capacity limit(s) for mental imagery representations per se and downstream computations.

### 4.3. Is “mental object” one or multiple natural kinds?

We demonstrated that mental objects created by linguistic instructions also drive posterior alpha suppression, similar to mental objects created by direct visual experience in recent past. This suggest that these two types of mental objects may be supported by the same system. Do all “objects” belong to the same ‘natural kind’ in this sense? This is not necessarily the case: for example, although phonological units (e.g., phonemes, syllables) have been seen as “auditory objects” (Griffiths & Warren, 2004), increasing the load of phonological units is accompanied by an increase of posterior alpha power (Proskovec et al., 2019) as opposed to a decrease. This may suggest that phonological units belong to another category of mental objects. Therefore, more studies with a variety of tasks and modalities need to be conducted in order to carve the joints of “mental objects”, i.e., what kind of mental objects belong to the same natural kind, and where to draw the lines among different natural kinds (Yu, 2024). This proposed line of research can thus also serve as an empirical reply to the long-lasting debate in the philosophy of mind, as to whether our brain has one vs. many symbolic systems (i.e., Language of Thoughts; Fodor, 1975; Mandelbaum et al., 2022), by focusing on the extent to which the same “mental objects” – the building blocks of symbolic computations in the mind – are truly shared across different tasks, modalities and neural systems.

## 5. Conclusion

We found that object-based load of mental objects derived from speech (a non-visual source) modulates left posterior alpha power, in a mental imagery task while participants’ eyes were closed. This pattern resembles what has been observed for visually-derived mental object load, where these mental objects are derived from direct visual input in recent history (e.g., in visual working memory delay-match-to-sample tasks). This suggests that object-based load for mental objects from difference sources may be accounted for by a shared neural resource. Taken together, our results support the hypothesis that mental objects tax a shared set of pointers in the mind, irrespective of their source modality.

## Data availability

Data and analysis scripts will be openly shared upon acceptance.

## Acknowledgements

We would like to thank Yifan Zeng, William Idsardi and Sebastián Mancha for discussions.

## Declare of interests

The authors declare no conflict of interests.

## Author contributions

Xinchi Yu: conceptualization, data curation, formal analysis, investigation, methodology, visualization, writing – original draft, writing – review & editing

Utku Türk: investigation, writing – review & editing

Allison Dods: investigation, writing – review & editing

Xing Tian: resources, supervision, writing – review & editing

Ellen Lau: conceptualization, funding acquisition, methodology, resources, supervision, writing – original draft, writing – review & editing

