## Supplementary Figure S1, S2 for "Non-visually-derived mental objects tax “visual” pointers"

### Supplementary Materials

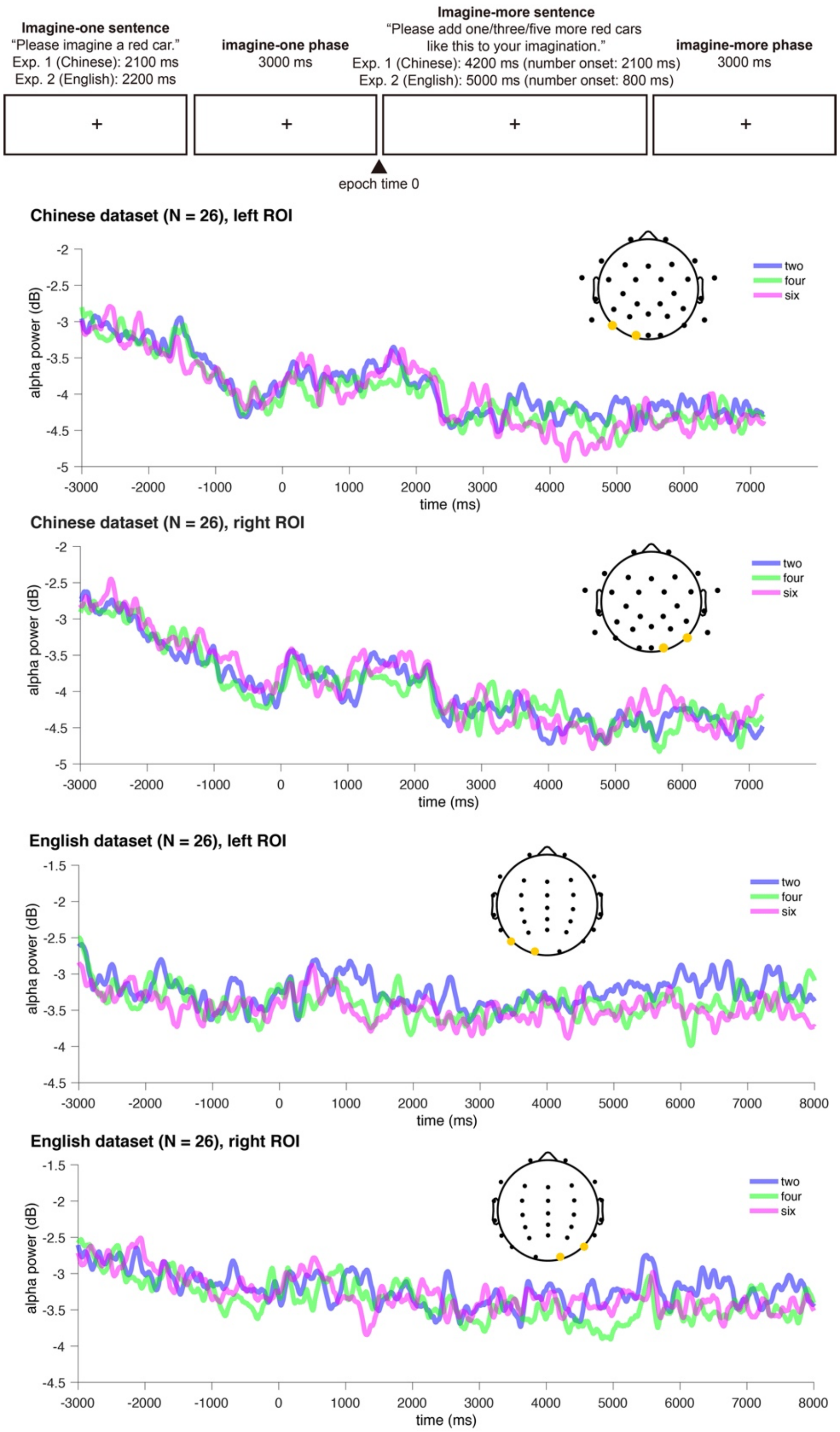


**Figure S1. Posterior alpha power in the left and right ROIs, illustrated across the full time period of the trial from the imagine-one phase to the end of the imaging-more phase.**

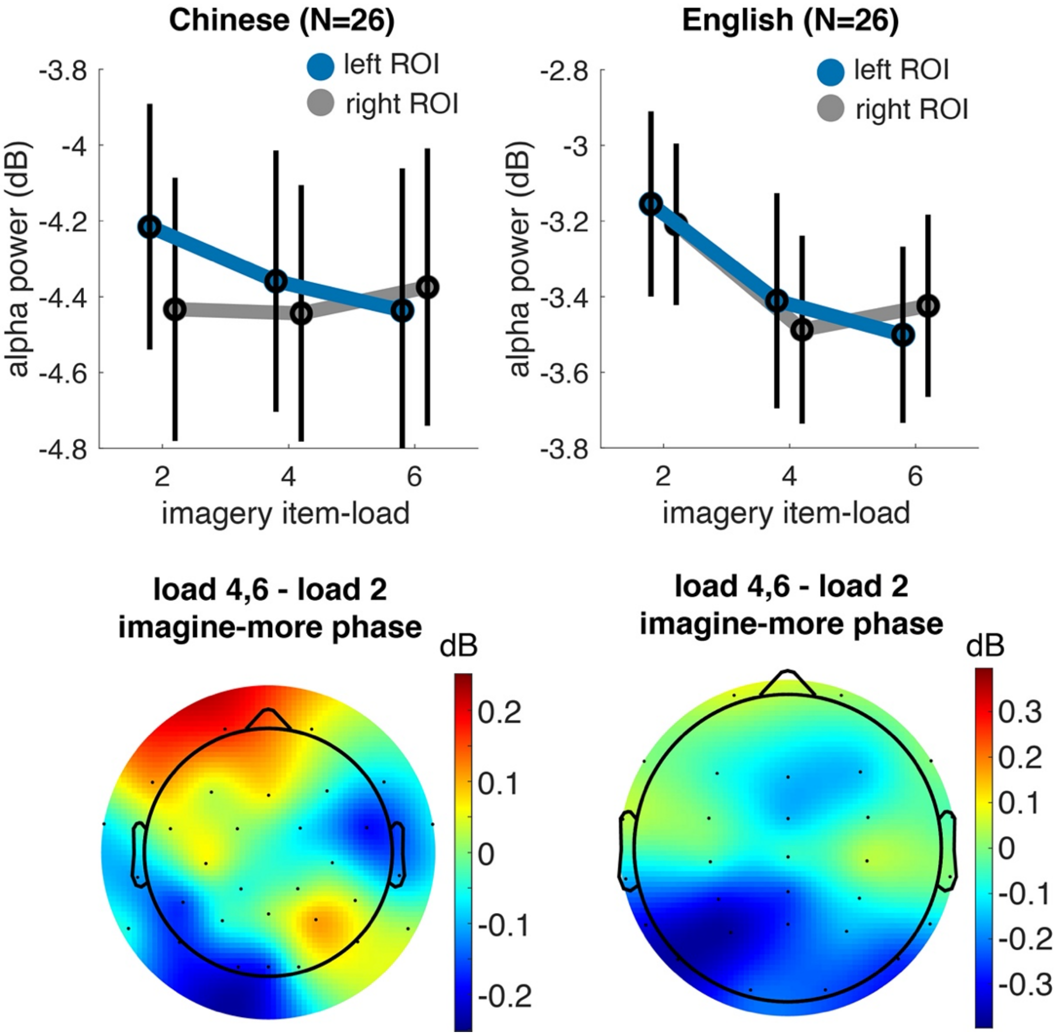


**Figure S2. Posterior alpha power during the imagine-more phase and scalp distribution for the English and Chinese participants respectively.** Error bars represent standard error across participants.
